# Computational Rational Design of Larger AAV Icosahedral Capsids

**DOI:** 10.64898/2026.05.09.724008

**Authors:** Michael D. Cioffi, Antoni Luque

**Affiliations:** Department of Biology, University of Miami, Coral Gables, FL, USA; Department of Physics, University of Miami, Coral Gables, FL, USA; Department of Chemical, Environmental and Materials Engineering, University of Miami, Coral Gables, FL, USA

**Keywords:** Adeno-associated virus (AAV), viral vectors, capsid engineering, molecular simulations, geometrical theory of viral capsids

## Abstract

Adeno-associated virus (AAV) is the preferred viral vector platform in gene therapy. Yet its packaging capacity, about 4.7 kb (kilobases), limits its therapeutic potential and represents a major bottleneck in the field. The packaging capacity of AAV is constrained by its small capsid, which forms a 26-nm-diameter shell assembled from 60 capsid proteins in a T=1 icosahedral architecture. Here, we propose increasing the cargo capacity of AAV vectors by engineering the next possible icosahedral architecture, T=3 (180 capsid proteins), which is predicted to provide a fivefold increase in volume capacity. Oligomers of VP3, the main capsid protein of AAV, were folded using AI-based methods. This identified triangular trimers as the optimal multimer compatible with the tiles of icosahedral lattices in the geometrical theory of capsids. The VP3 trimers were assembled into a T=3 architecture and coarse-grained at 5Å resolution. It was necessary to introduce 15 deletions (VP3Δ15) to accommodate the T=3 curvature. Molecular simulations under physiological conditions demonstrated the stability of the 45 nm-diameter T=3 capsid. Structural analysis measured a five- to sixfold increase in internal volume and estimated a potential upper cargo limit of 35 kb. The engineered VP3Δ15 could enable delivery of multicistronic constructs, larger regulatory elements, and CRISPR systems beyond the reach of current AAV vectors. Additionally, the introduced generalized protein design framework could be used to engineer capsids with larger T-numbers and to modify the capacity of other icosahedral delivery systems.

## INTRODUCTION

Adeno-associated virus (AAV) is a small, single-stranded (ss) DNA virus that infects humans and has a genome of approximately 4.7 kilobases (kb) (Agbandje-McKenna & Kleinschmidt, 2012; Zengel & Carette, 2020). AAV belongs to the *Parvoviridae* family and encodes the two characteristic gene cassettes or open reading frames (Kazlauskas et al., 2019). Rep regulates replication, and Cap encodes the structural proteins (Figure 1a). Unlike other human viruses in this family, AAV requires a helper virus, such as adenovirus, human papillomavirus, or herpesvirus, to replicate, as reflected in its genus name, *Dependoparvovirus* (Cotmore et al., 2019). AAV exhibits broad tropism across its 13 serotypes (AAV1–AAV13) but is not known to cause disease in humans (D. Wang et al., 2019; J.-H. Wang et al., 2024). The cloning of AAV enabled the development of recombinant AAV vectors, which exploit AAV’s broad cellular tropism, lack of known pathogenicity, and ability to persist primarily as episomal DNA to support long-term transgene expression (Samulski et al., 1982, 1989; D. Wang et al., 2019; J.-H. Wang et al., 2024). In recombinant AAV vectors, the viral rep and cap coding regions are removed from between the inverted terminal repeats and replaced with a transgene expression cassette, while Rep and Cap functions are supplied in trans during vector production (Muzyczka, 1992; Samulski et al., 1989). AAV vectors have become a leading platform for gene therapy due to their low pathogenicity, minimal cytotoxicity, and broad tissue tropism, enabling long-term, stable gene expression across diverse cell types (Gray et al., 2010; Suarez-Amaran et al., 2025; J.-H. Wang et al., 2024). The 2017 approval of Luxturna by the U.S. Food and Drug Administration marked a major milestone for AAV-based therapeutics (Russell et al., 2017; Smalley, 2017). Yet the therapeutic use of AAV vectors is constrained by their limited genomic capacity (<5 kb) (Agbandje-McKenna & Kleinschmidt, 2012; Zengel & Carette, 2020).

**Figure 1.**
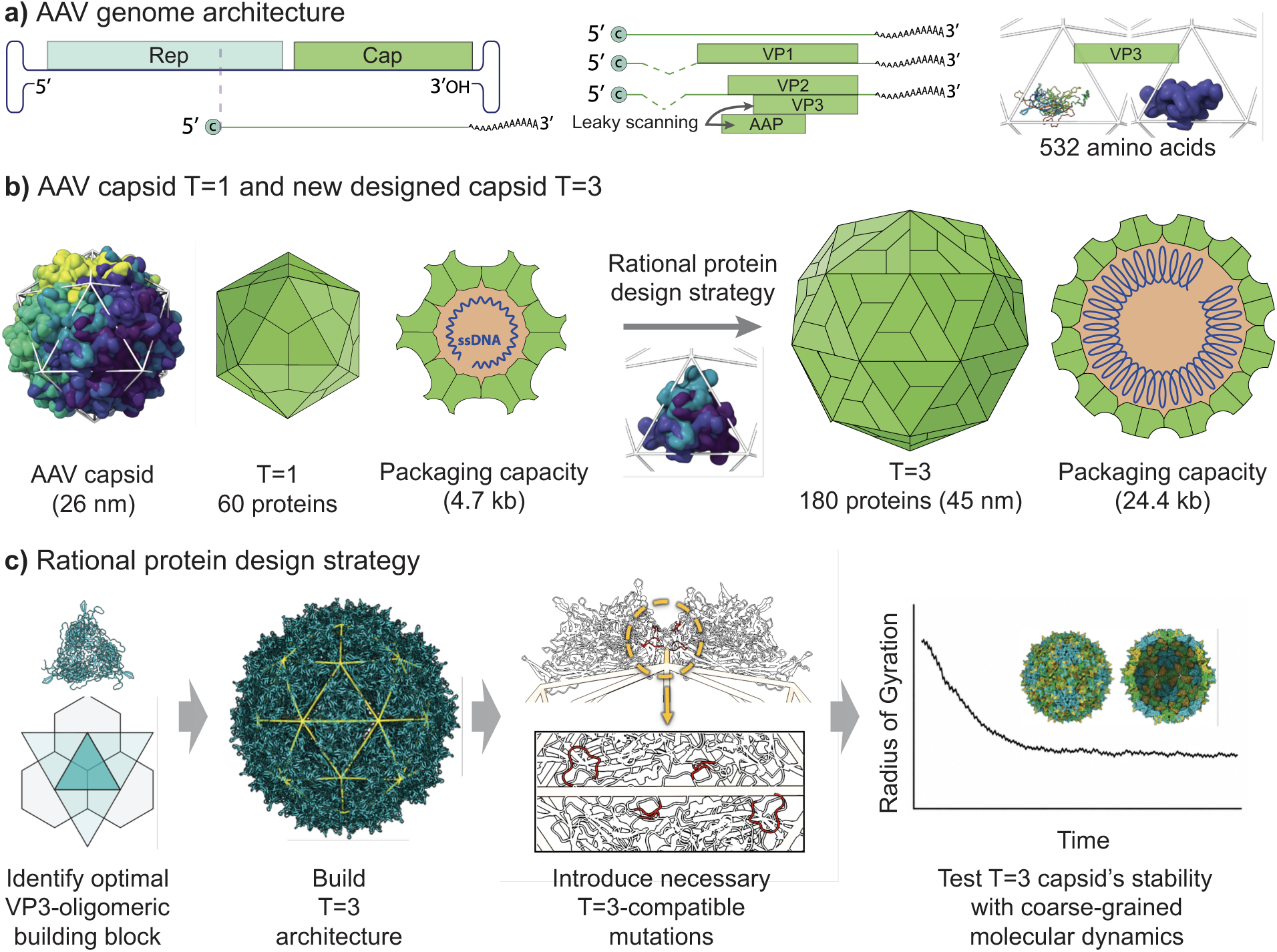
AAV capsid and strategy to enlarge its capacity. **a)** Architecture of the AAV genome and capsid proteins encoded in the Cap open reading frame (left and middle panels). The structure and amino acid length of VP3 is displayed (right panel). **b)** Canonical capsid formed by AAV (panels on the left), displaying the diameter, T-number architecture, number of capsid proteins, and packaging capacity. Target capsid and expected packaging capacity (panels on the right). **c)** Key steps implemented in the rational protein design strategy. See text in the introduction and result figures 2-4 for details. Panels **a)-b)** adapted vectorial images from ViralZone (De Castro et al., 2024) and molecular renders ofthe PDB 1LP3 structure (Xie et al., 2002) generated in ChimeraX (Meng et al., 2023).

The limited packaging capacity of AAV restricts its spectrum of treatable diseases (D. Wang et al., 2019). AAV vectors can deliver diverse genetic payloads, including cDNA expression cassettes, shRNAs, CRISPR components, and gene-editing nucleases, but their packaging capacity is limited to approximately 4.7–5.0 kb, including ITRs and regulatory elements (Dong et al., 1996; Naso et al., 2017; J.-H. Wang et al., 2024). This constraint limits single-vector AAV delivery for large-gene replacement therapies and some genome-editing approaches, particularly for rare genetic diseases involving large therapeutic transgenes that exceed the AAV packaging limit (Ferreira, 2019; Greene et al., 2023; Zhou et al., 2024). Examples include ABCA4 in Stargardt disease (Auricchio et al., 2015), F8/Factor VIII in hemophilia A (High & Anguela, 2016), and DMD/dystrophin in Duchenne muscular dystrophy (Lostal et al., 2014; Rodino-Klapac et al., 2007), which require coding sequences of approximately 6.8 kb, 7.0 kb, and >11 kb, respectively. The field is pursuing several strategies to overcome this limitation. One strategy for overcoming AAV genome-size constraints is the use of miniaturized transgenes, including mini-/micro-dystrophins, combined with sequence-level optimization to improve expression efficiency (Chwalenia et al., 2025; Foster et al., 2008; Piepho et al., 2023). A more promising approach is the use of dual AAV vectors (split AAV), which can double the cargo delivered to a target cell but requires co-infection with different AAV vectors, reducing efficacy and increasing toxicity (Ertl, 2022; McClements & MacLaren, 2017; Trapani et al., 2014). The optimization and dual-vector strategies emerged because the most straightforward strategy of increasing the genetic cargo of a single AAV vector has faced important limitations, as even modest genome expansion, nearly or above 5 kb, reduces transduction efficiency (Suarez-Amaran et al., 2025; Z. Wu et al., 2010). This is due to a physical constraint: the viral genome occupies a significant fraction of the internal volume of the capsid (Luque et al., 2020; Sun et al., 2017), and increasing the genome content impacts the capsid stability and the conformational changes needed for genome delivery (Horowitz et al., 2013).

The reduced internal volume in AAV vectors is associated with the small dimensions of its capsids, which form a 26-nm-diameter shell assembled from 60 capsid proteins arranged in a T=1 icosahedral architecture (Agbandje-McKenna & Kleinschmidt, 2012; Xie et al., 2002) (Figure 1b). The prevalence of T=1 across *Parvoviridae* viruses(Cotmore et al., 2019; Mietzsch et al., 2019), which means “small” or “tiny” in Latin, may explain why mutagenesis efforts on the capsid protein have successfully altered AAV tropism, antigenicity, and transduction efficiency (Büning & Srivastava, 2019; Pupo et al., 2022; P. Wu et al., 2000). Yet, to our knowledge, there has been no characterization of a capsid larger than T=1.

The triangulation number (T) quantifies the number of major capsid proteins (or quasi-equivalence positions) in the asymmetric unit of icosahedral capsids and is proportional to the number of major capsid proteins, 60T (Caspar & Klug, 1962; Twarock & Luque, 2019). Icosahedral symmetry restricts the possible values of the T-number, which follow a precise sequence, with T=1, 3, 4, 7 as the first elements (Figure 1b). Viral capsids in other viral families have been reported to adopt a vast range of icosahedral architectures, from T=1 to T>1000 for giant viruses (Luque et al., 2020; Mietzsch et al., 2025; Suhanovsky & Teschke, 2015; Xiao & Rossmann, 2011). This includes viruses that can form both T=1 and T=3, such as brome mosaic virus (BMV) (Larson et al., 2005; Lucas et al., 2002) and cowpea chlorotic mottle virus (CCMV) (Speir et al., 1995), whose capsid proteins share a similar structural fold, the jelly-roll fold, observed in AAV capsid proteins (Figure 1b). The geometrical theory of viral capsids predicts that, for a given major capsid protein, a T=3 capsid provides a fivefold increase in internal volume, V/V_0_ = (T/T_0_)^3/2^ = (3/1)^3/2^ = 5.2 (Lee et al., 2022; Luque et al., 2020; Twarock & Luque, 2019). Thus, regardless of the evolutionary reason why AAV and other parvoviruses do not form capsids larger than T=1, the possibility of increasing the capsid architecture of AAV to form T=3 capsids could increase its packaging capacity to around 30 kb (Figure 1b), providing a major expansion to the applicability of AAV-based therapeutics.

Here, we addressed this problem computationally by engineering the AAV capsid protein VP3 to form stable T=3 capsids (Figure 1b). AAV’s structural cassette, Cap, encodes three overlapping capsid proteins, VP1, VP2, and VP3, that share a 533-residue core corresponding to VP3 (Oyama et al., 2021; Wörner et al., 2021). VP3 is the main component of the AAV capsid, with an average stoichiometry of 10:1:1 relative to VP1 and VP2 (Wörner et al., 2021). *In vitro* experiments have demonstrated that VP3 can assemble into VP3-only virus-like particles in the absence of VP1 and VP2 (Mietzsch et al., 2023; Sonntag et al., 2011). The sequence of VP3 was folded using AI-based methods to identify the optimal VP3 multimers compatible with the geometrical tiles in the icosahedral theory of viral capsids (Twarock & Luque, 2019) (Figure 1c). This optimal tile was used to generate T=1 and T=3 architectures and analyzed to identify amino acid regions that required modifications to accommodate the new larger architecture. Coarse-grained molecular dynamics (CGMD) simulations were performed to assess the stability of the new structure (Figure 1c), since viral capsid architecture is governed by a balance between geometric constraints, inter-subunit interactions, and collective dynamical behavior, which together determine physical stability, assembly, and mechanical response (Luque & Reguera, 2024; Roos et al., 2010). Structural analysis confirmed the stability of the T=3 capsids and estimated up to a 6-fold increase in the internal packaging capacity. These results described and discussed in detail below provide an exciting parent mutant for the field to explore enlarged AAV vectors to address one of the major bottlenecks in the field. The engineering effort focused on the AAV2 serotype because its capsid is one of the most extensively structurally characterized AAV systems (Agbandje-McKenna & Kleinschmidt, 2012; Drouin & Agbandje-McKenna, 2013). But the mutations engineered can be adapted to other serotypes. Besides, the generality of the computational design approach opens also the doors to design other T-numbers and modify the architecture of other delivery systems of interest.

## RESULTS

### Triangular VP3 trimers were the optimal icosahedral building block for AAV2 capsid design

The optimal tiles compatible with icosahedral lattices for AAV2 capsids were identified by folding VP3 oligomers using AlphaFold 3 (AF3) (Abramson et al., 2024). The VP3_n_ trimer (n=3), VP3_3_, yielded an interface predicted template modeling (ipTM) score of 0.75, significantly higher than the ipTM scores of all other oligomers folded, which yielded below 0.5 (Figure 2a). VP3_3_ displayed a quasi-regular triangular shape (Figure 2a). None of the other oligomers formed symmetrical structures except the dimer (n=2). The predicted template modeling (pTM) score for the VP3 trimer was 0.81, the only oligomer to score above the high-quality threshold (pTM > 0.75), and it trailed only the VP3 monomer (pTM = 0.93). The high pTM and ipTM scores for the VP3 trimer are consistent with observations that VP trimers are preferential oligomers in AAV capsid assembly (Maurer et al., 2018). Therefore, our optimal oligomer, VP3_n*_, was selected to be the trimer (n*=3).

**Figure 2.**
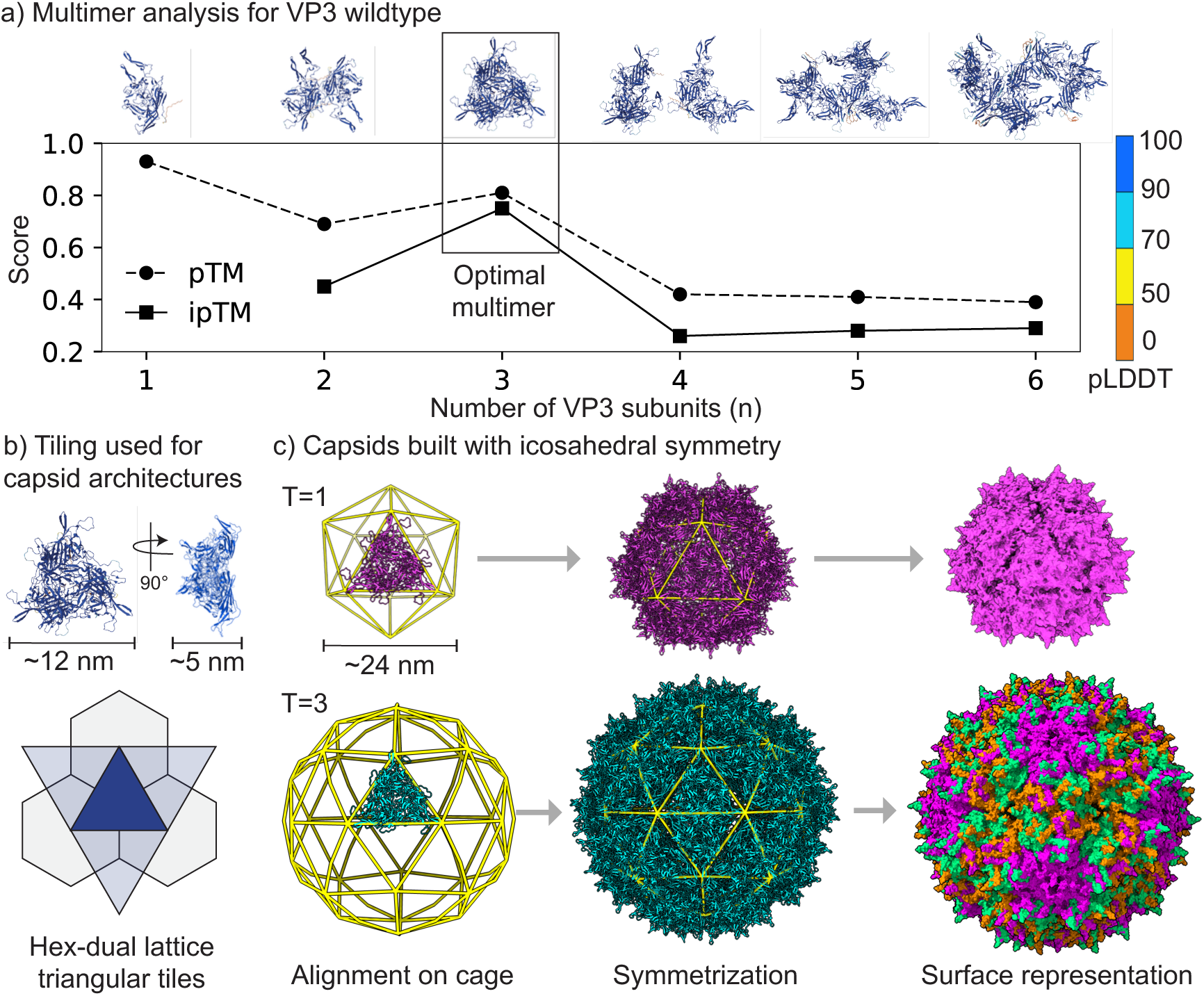
Optimal building blocks in AAV2. **a)** AlphaFold 3 ipTM (solid line) and pTM (dashed line) scores for the folded AAV2 VP3 monomer and oligomers built with up to six subunits (n). The folded structures are displayed above the plot. The color coding corresponds to the pLDDT score. Full size images of each oligomer have been added to Figure S1, where local scores are easier to interpret. The black box highlights the optimal oligomer selected for the icosahedral design. b) The selected optimal oligomer (triangular VP3 trimer) is shown above the icosahedral lattice that is congruent with the oligomer’s shape. The same color (turquoise) is used for the oligomer and the tile’s lattice to facilitate their comparison. c) T=1 icosahedral cage displaying the initial placement of the triangular VP3 trimer (ribbon style), the cage and built capsid (ribbon style), and surface-style representation of the full capsid. T=3 cage and capsid following the same logic. The proteins in surface representation for the T=3 structure are displayed in three different colors, corresponding to the three quasi-equivalent positions of the T=3 capsid.

The regular triangular shape of the VP3_3_, with side lengths of ∼12 nm and a thickness of ∼5.6 nm (Figure 2b), was congruent to the triangular geometrical tiles that are associated with the hexagonal-dual icosahedral lattice (Figure 2b). Based on the geometrical theory of viral capsids (Twarock & Luque, 2019), this regular triangular trimer satisfies the geometrical requirements to build potentially any T-number icosahedral capsid. Thus, T=1 and T=3 capsids (Figure 2c) were built using folded VP3 trimers (see Methods for details). These initial capsid models displayed approximate diameters of ∼30 nm and ∼41 nm, respectively.

### T=1 VP3_3_ capsid is stable while T=3 VP3_3_ is indeterminate

To validate our method of capsid construction based on optimized icosahedral tiling, coarse-grained molecular dynamics simulations (∼5 Å resolution) were performed with the VP3_3_ VP3 T=1 capsid and compared to the experimentally resolved AAV2 T=1 capsid derived from X-ray crystallography (Xie et al., 2002) (Figure 3a). Trajectory analysis over 2 μs revealed global structural stability for both systems, as indicated by the radius of gyration (*R*_g_) and radial measurements; both exhibited minimal fluctuations throughout the simulation period (Figure 3a). Windowed *R*_g_slope analysis showed that the AAV2 T=1 and VP3_3_ T=1 capsids quickly reached a low-drift regime, with |m| falling below the 0.25 nm/µs stability threshold within the 1st (0-0.1 µs) and 6th (0.5-0.6 µs) windows, respectively (Figure S2). The average *R*_g_ in the stable region was calculated to be 11.0±0.25 nm for the T=1 AAV2 PDB structure and 11.5±0.27 nm for the T=1 VP3_3_ structure. Average radial measurements of the two T=1 capsids were calculated for the same time range. The average minimum radii are 7.8±0.25 nm (PDB) and 7.9±0.25 nm (VP3_3_), average mid-radii are 11.0±0.25 nm (PDB) and 11.4±0.25 nm (VP3_3_), and average external radii are 15.1±0.25 (PDB) and 15.9±0.25 nm (VP3_3_). The *R*_g_ and R_avg_ values for each system are well within the uncertainty reflecting a highly spherical structure. The VP3_3_–assembled capsid exhibited slightly higher maximum radii compared to the PDB-derived structure, while maintaining an essentially identical minimum and average radius (within model resolution), indicating that the inner shell boundary is conserved. This behavior is consistent with localized deviations in inter-subunit geometry rather than a uniform expansion of the capsid since R_min_ stays constant. In the VP3_3_ model, small differences in trimer tiling—such as minor rotational offsets, altered interfacial angles, or less optimal packing at quasi-equivalent positions—are more prone to produce subtle outward displacements in specific regions of the shell due to the capsid building procedure described in the methods. These effects are most pronounced at symmetry interfaces, where small perturbations in local curvature can increase the radial extent of a subset of residues or beads. As a result, the capsid remains structurally stable but exhibits a slightly broadened radius distribution, reflecting minor surface irregularities and reduced symmetry enforcement relative to the PDB-derived reference. Root mean square deviation (RMSD) analysis relative to the initial structures showed a gradual increase in both systems, but the values remained below the model resolution threshold of 0.5 nm throughout the trajectory (Figure S3). Rather than reaching a strict plateau, the RMSD profiles exhibited a semi-plateau, which is expected for capsid simulations because inter-subunit interfaces remain dynamically active. Continuous small-scale rearrangements at these interfaces can produce slow residual drift, even in structurally stable assemblies. These combined results show that the VP3_3_ assembly protocol produces capsids with consistent structural profiles that differ by less than model resolution in R_g_ and R_avg_, thereby validating the methodology for subsequent applications to non-native capsid architectures.

**Figure 3.**
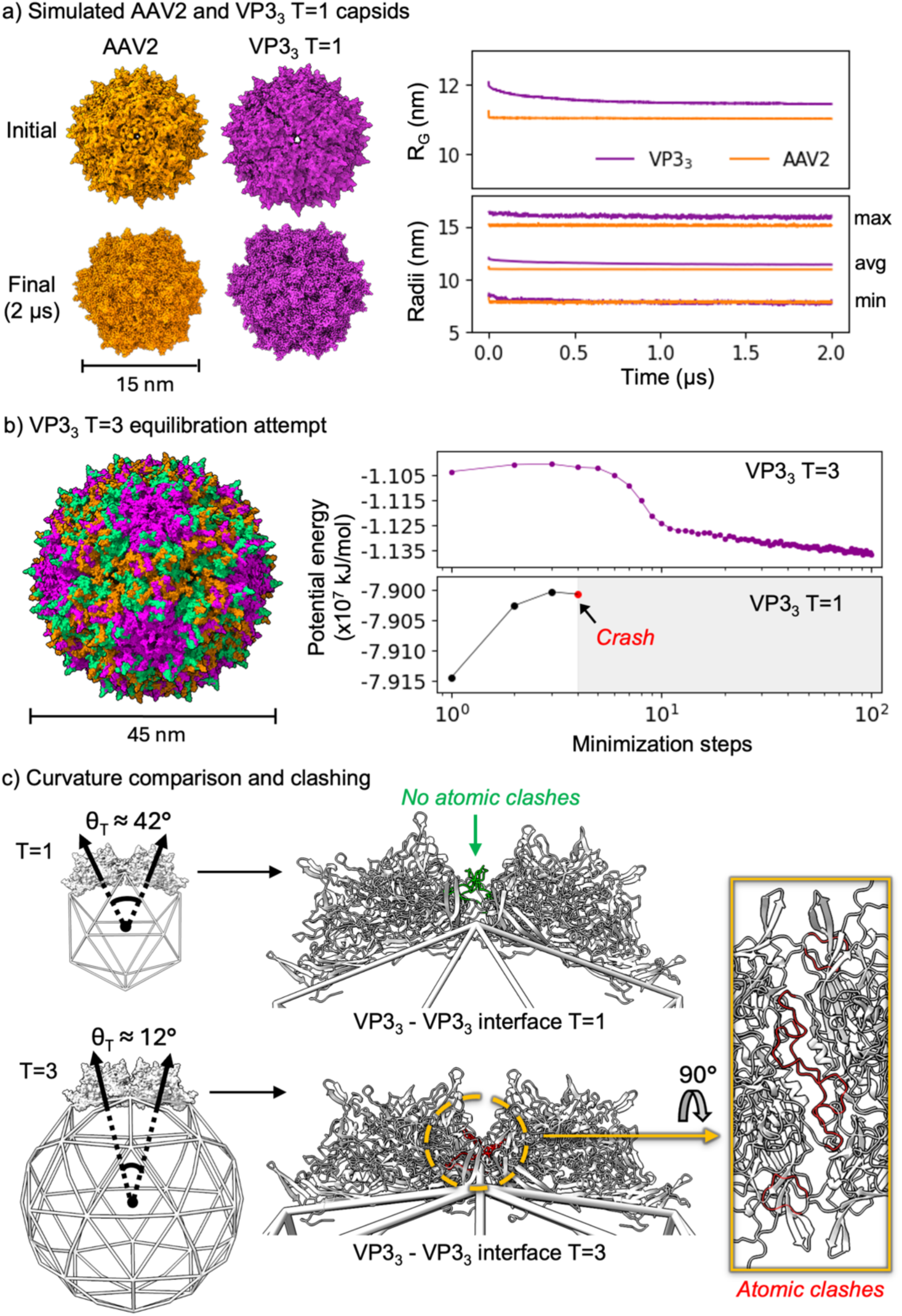
Stability analysis of VP3_3_ T=1 and T=3 capsids. **a)** Initial (top) and final (bottom) capsid structures for the coarse-grained models of VP3_3_ AAV2 T=1 PDB (orange) and VP3_3_ (magenta). The two right panels plot the radius of gyration (*R*_g_) and the minimum (R_min_), average (R_avg_), and maximum (R_max_) radii throughout the molecular dynamics (MD) trajectory. b) The initial VP3_3_ T=3 coarse-grained model, where proteins are represented using three colors (orange, lime, magenta), based on their quasi-equivalent positions in the icosahedral capsid. The right panel plots the potential energy of this capsid in the initial (MD) equilibration steps (black dots joined by a solid line). The red dot indicates the step where equilibration could not progress (crash). The potential energy per step for the successfully equilibrated VP3_3_ T=1 capsid (magenta line) is provided for contrast. c) Comparison of facet-to-facet curvature (left), where normal vectors are displayed for adjacent trimeric tiles. The angle θ_*T*_ denotes the approximate angle between the normals of neighboring tiles, with θ_*T*_ ≈ 12^∘^for the T=3 assembly and θ_*T*_ ≈ 42^∘^for the T=1 assembly. The molecular VP3_3_-VP3_3_ interface is shown for the initial VP3_3_ T=1 and VP3_3_ T=3 capsid structures (middle), highlighting entire loop regions 656–667 and 695–718. These regions are colored green in the T=1 structure to indicate non-clashing and red in the T=3 structure to indicate the presence of steric clashes. A close-up view of the loop regions in the VP3_3_-VP3_3_ T=3 interface is shown on the right, with the same clash-prone regions highlighted in red. The PDB AAV2 T=1 capsid system contained 900,483 total CG beads, including 798,051 water, 2,946 Na⁺, and 2,826 Cl⁻ beads. The VP3_3_ T=1 capsid system contained 955,105 total CG beads, including 850,233 water, 3,236 Na⁺, and 2,996 Cl⁻ beads. The VP3_3_ T=3 capsid system contained 6,981,346 total CG beads, including 6,642,612 water, 24,557 Na⁺, and 24,197 Cl⁻ beads.

The next question was whether AAV2 VP3_3_ could stabilize in T=3 capsid architecture in the absence of any sequence modifications. Multiple variations of the initial trimer placement in the VP3_3_ T=3 configuration were attempted, but all produced similar failures during the equilibration stage of the simulation (Figure 3b). During the initial equilibration time steps, the total potential energy increased rapidly by (∼0.015 x10^7^ kJ/mol). before the simulation terminated. This rapid energy increase and immediate equilibration failure suggest that, although the structure could be minimized, residual steric strain remained at the trimer–trimer interface and became dynamically unstable once thermal motion was introduced. In contrast, the VP3_3_ T=1 capsid displayed only a minor initial increase in potential energy (∼0.005 kJ/mol), followed by a sharp decrease (∼0.020 x10^7^kJ/mol), and stabilization at ∼0.032 x10^7^ kJ/mol below its initial value (Figure 3b). The inability of the VP3_3_ T=3 capsid to remain stable during equilibration was associated with protein clashes that generated prohibitive van der Waals repulsion energies at the trimer–trimer interface (Figure 3c). Ultimately, this was caused by the dihedral angle at trimer-trimer interfaces, which increased by ∼22% in the T=3 architecture (angle≈168°) compared to the T=1 architecture (angle≈138°) (Figure 3c). The corresponding normal–normal angle between adjacent trimeric tiles is ∼12° in the T=3 assembly and ∼42° in the T=1 assembly, indicating that neighboring tiles are locally flatter in the larger T=3 architecture. This reduction in curvature caused two flexible loop regions at the trimer-trimer interface, residues ∼656-667 (VP1) and ∼695-718 (VP1), to interpenetrate beyond sterically permissible distances in the T=3 architecture (Figure 3c). Given that the T=3 architecture has 60 trimer-trimer interfaces, compared to 20 in the T=1 architecture, the cumulative energetic penalty prevented the successful equilibration of the T=3 capsid. Thus, wild-type AAV2 VP3_3_ could not be evaluated for stability in a T=3 architecture *in silico*.

### Deletion of clashing amino acids yields stable T=3 capsids

It was hypothesized that if residues involved in the clashes at the VP3 trimer-trimer interfaces were removed then the mutant, VP3Δ, would yield stable T=3 capsids. Deletion sites were chosen to reduce interfacial clashes while retaining residues expected to support capsid stability, including hydrophobic residues (Pace et al., 1996, 2011), and avoiding positions previously identified as critical for capsid assembly or function (P. Wu et al., 2000) (Table S1). This led to a VP3 mutant sequence containing 15 deletions, VP3Δ15 (Figure 4a). Folding the modified sequence confirmed that triangular trimers, VP3Δ15_3_, were the optimal oligomer as in VP3_3_, but this time they also yielded folded trimer-trimer structures, VP3Δ15_3_-VP3Δ15_3_, for 6-mers (Figure S1). This trimer-trimer structure did not form for VP3, suggesting that the deletions are particularly adept at forming trimer-trimer interfaces.

**Figure 4.**
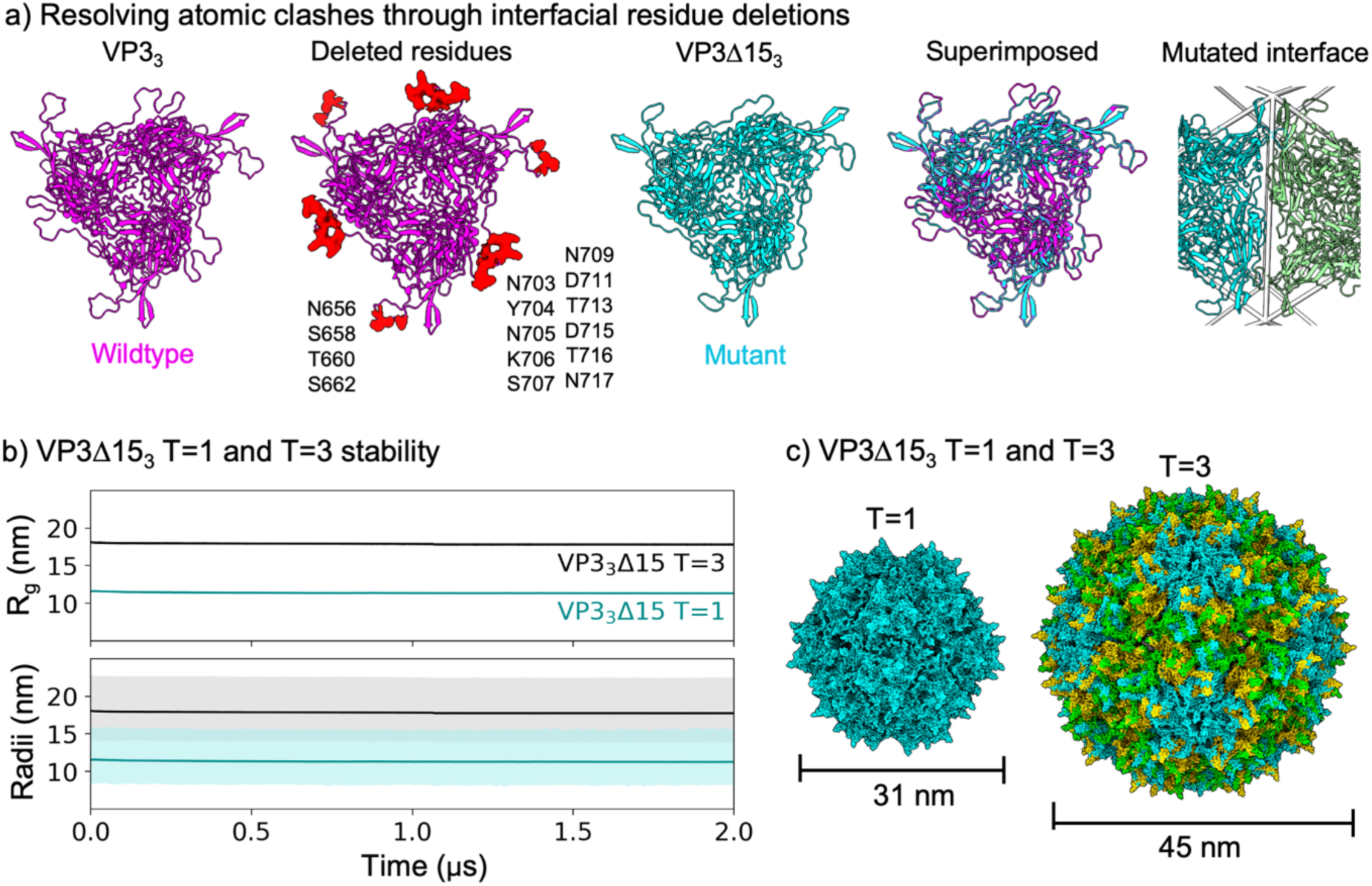
Stability of VP3Δ15_3_ T=3 capsids. a) Comparison of the WT VP3 trimer (magenta) and the mutated VP3Δ15 trimer (cyan). The amino-acid and sequence location of each deletion (in red) is displayed (second panel to the left). The number is the location in the VP1 reference sequence, which contains VP3. The panel on the right displays the trimer-trimer interface of two mutated trimers in cyan and green, respectively. b) Time evolution of the radius of gyration and radii components for VP3Δ15_3_ T=1 (cyan) and VP3Δ15_3_ T=3 (black) for the 2 µs trajectories. The light semi-transparent shaded regions indicate the extent of the maximum and minimum radii measurements for each respective system. c) Structural comparison of the end points for the VP3Δ15_3_ T=1 and VP3Δ15_3_ T=3 capsids. The proteins are colored based on their quasi-equivalent positions: one color (cyan) for T=1 and three colors (cyan, green, and yellow) for T=3. The VP3Δ15 T=1 capsid system contained 914,456 total CG beads, including 815,460 water, 2,938 Na⁺, and 2,878 Cl⁻ beads. The VP3Δ15 T=3 contained 4,527,123 total CG beads, including 4,216,635 water, 15,564 Na⁺, and 15,384 Cl⁻ beads.

Both VP3Δ15_3_ T=1 and VP3Δ15_3_ T=3 architectures were simulated for 2 µs (Figure 4b). This duration ensured adequate sampling of capsid stability and dynamic behavior (see Methods for justification). Windowed *R*_g_ slope analysis showed that the VP3_3_Δ15 T=1 and VP3_3_Δ15 T=3 capsids reached a low-drift regime, with |m| falling below the 0.25 nm/µs stability threshold within the 5th (0.4-0.5 µs) and 11th windows (1-1.1 µs), respectively (Figure S2). The VP3_3_Δ15 T=3 capsid displayed the latest stabilization time, consistent with its larger size and correspondingly slower global relaxation dynamics. During the stable period, the radius of gyration, *R*_g_, of VP3Δ15_3_ T=1 was 11.3±0.25 nm, while VP3Δ15_3_ T=3 had an average *R*_g_ of 17.8±0.25 nm from 1 µs to the end of the simulation period (Figure 4b). Both gyration radii were well within the uncertainty of the average radii (VP3Δ15_3_ T=1: ∼11.30±0.28 nm; VP3Δ15_3_ T=3: ∼17.76±0.25 nm), indicating high sphericity. This combination of *R*_g_ and Radii measurements confirm the hypotheses proposed in that VP3Δ15_3_ T=3 yields a stable capsid. However, VP3Δ15_3_ also yielded a stable T=1 capsid, indicating that this smaller structure could compete with T=3 in the assembly process.

The geometrical theory of viral capsids predicts that, assuming a similar sphericity, a T=3 would display a radius 73%, (T/T0)^1/2^ ≈ 1.73, larger than a T=1 capsid assembled from the same capsid protein. The average radius of VP3Δ15_3_ T=1 was 11.30±0.28 nm and the average radius of VP3Δ15_3_ T=3 was 17.76±0.25 nm, yielding a 1.58±0.05 times increase in radius for T=3, slightly below the predicted theoretical value. This reduced scaling factor reflects deviations from strict geometric similarity in the assembled molecular shells during our capsid-building protocol, including differences in subunit tilt and interface organization, changes in the effective surface area occupied per subunit, initial symmetrization of VP3_3_ models, and structural relaxation during coarse-grained equilibration, all of which can compact the T=3 shell relative to the ideal Caspar–Klug prediction. The analysis of the root mean squared deviation (RMSD) with the initial structures revealed an overall increase in both cases which remained below the model resolution for the entire trajectory, < 0.5 nm (Figure S3). A semi-plateau in RMSD, rather than a strict plateau, is expected in capsid simulations due to the dynamic nature of inter-subunit interactions. These interfaces undergo continuous, small-scale rearrangements over time, resulting in a slow drift rather than complete structural convergence.

### The VP3Δ15_3_ T=3 capsid displays over five times more packaging capacity than AAV T=1

The main rationale for designing T=3 capsids was the increase in the potential packaging capacity of AAV vectors. The working hypothesis, based on the geometrical theory of viral capsids, was that a T=3 capsid could offer an increase of more than five times in volume, V, compared to a T=1 capsid, V/V_0_ = (T/T0)^3/2^ ≈ 5.2. This theoretical prediction was tested computationally by comparing the molecular simulation of AAV2 T=1 capsid, which was modeled based on the experimental reconstruction of a wild-type AAV2 capsid (PDB 1LP3), to the engineered VP3Δ15_3_ T=3 capsid (Figure 5).

**Figure 5.**
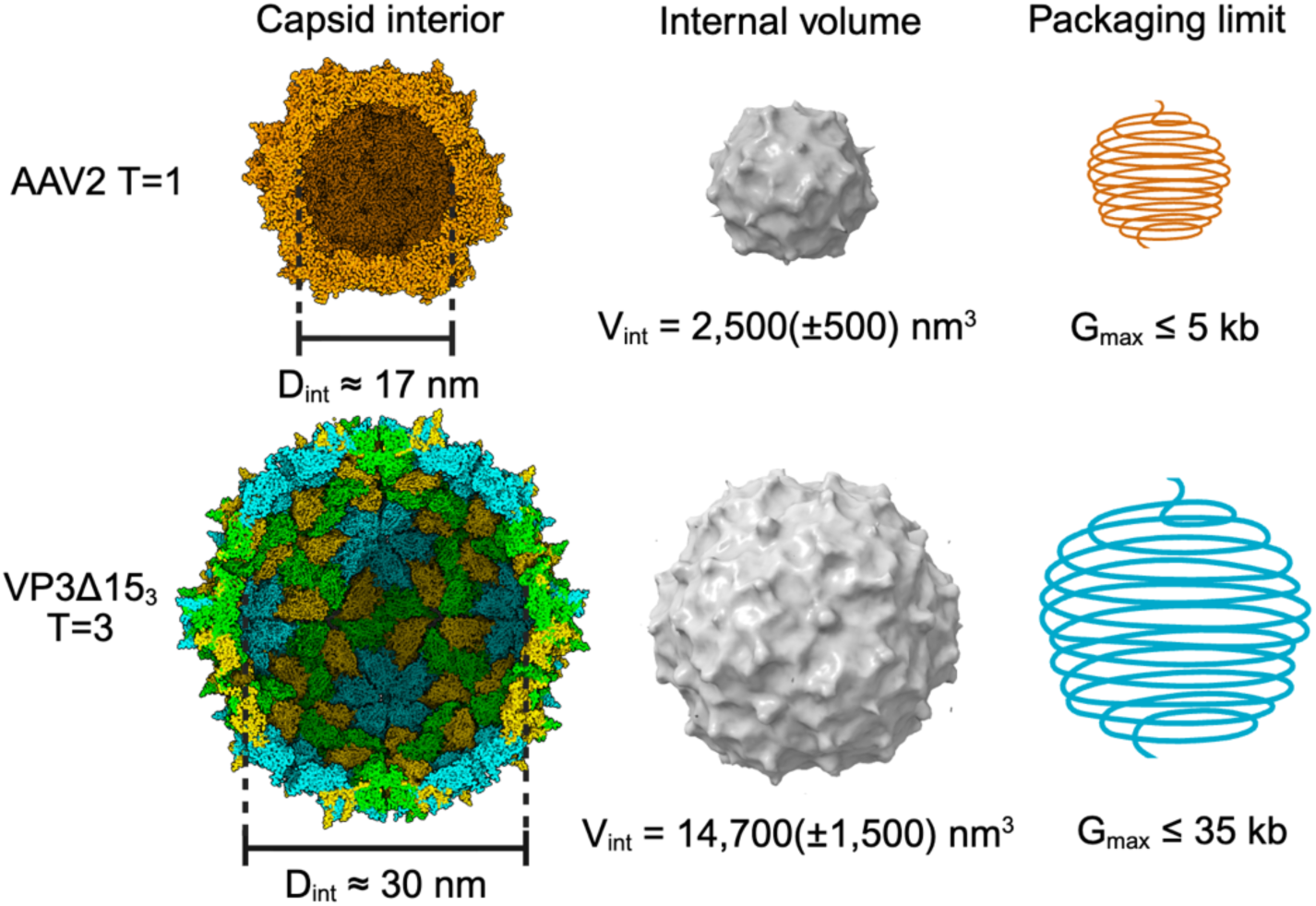
Estimated packaging capacity for VP3Δ15_3_ T=3. Rendering of the interior of the WT T=1 capsid (orange), its interior volume (grey), and a coiled polymer representing maximum genome capacity (orange). Rendering of the interior of the VP3Δ15_3_ T=3 capsid (tri-colored), its interior volume (grey), and a coiled polymer representing maximum genome capacity (cyan). The internal diameter (D_int_) of each capsid is shown on the left. The estimated value of the internal volume (V_int_) with error (δV) is displayed in the middle column. The estimated maximum genome capacity (G_max_) in kilobases (kb) is displayed for both the T=1 and T=3 capsids below the coiled polymers. G_max_ for the T=1 capsid was determined as the upper bound of empirical measurements. G_max_ for the T=3 capsid was calculated using a packaging density (ρ) of 1.92 ± 0.54 nt/nm^3^, which was derived from the internal volume calculations and empirical genome packaging capacity of AAV2 T=1.

A conservative estimate of the internal volume was obtained using the Blob tool in ChimeraX (Meng et al., 2023), which computes the volume enclosed by a surface. For both the PDB AAV2 T=1 and VP3Δ15_3_ T=3 capsids, the average interior volume (V_int_) was calculated from five late trajectory frames (1.90 µs to 1.98 µs, Δt=0.2 µs). The total uncertainty in volume, δV_total_ (referred to as δV), is dominated by the blob resolution, which was set to 1.5 nm to fill surface pores and eliminate holes in the capsid shell, thereby allowing the software to distinguish the internal and external regions. Values for δV were calculated as described in the methods. For the AAV2 T=1, we obtain a V_int_ (±δV) of 2,500 (±500) nm^3^, and for VP3Δ15_3_ T=3, we obtain a V_int_ (±δV) of 14,700 (±1,500) nm^3^.

Using these volume measurements, we are then able to compute an estimated genome packaging capacity for the simulated VP3Δ15_3_ T=3 capsid. Wild-type AAV2 was treated as having a single-stranded DNA genome size (G±δG) of approximately 4.8 ± 0.2 kb, with the rationale for this value described in the methods. Using this genome size and the measured V_int_ (±δV) of the simulated AAV2 T=1 capsid, the estimated packaging density (ρ±δρ) of the genome was calculated to be ∼1.9 ± 0.4 nt/nm^3^, which is in the range of known *Parvovirdae* packaging densities (1.5-3.0 nt/nm^3^) (Mietzsch et al., 2025). Then, from ρ and the measured V_int_ (±δV) of the simulated VP3Δ15_3_ T=3 capsid, we estimated the total genome capacity (G±δG) of VP3Δ15_3_ T=3 to be ∼28.2 (±6.5) kb. If we take the upper bound of the uncertainty in both G±δG values, we obtain a G_max_ = 5 kb for AAV2 T=1 and a G_max_ = 34.7 (≈ 35 kb) for VP3Δ15_3_ T=3. The complete workflow, including all equations and parameters, is described in detail in the Methods. The volume and packaging capacity derived from the simulations were compared with the expected values from the scaling laws of the geometrical theory of viral capsids. The internal volume of the simulated VP3Δ15_3_ T=3 structure is ∼5.9 (±1.4) times larger than that in the simulated WT T=1. The theoretical scaling value calculated from geometric theory (V/V_0_ = (T/T_0_)^3/2^ ≈ 5.2) falls well within the uncertainty range (4.5-7.3x) of the estimated volume increase as determined from the simulated structures. Even on the lower bound (4.5x), this is substantially larger than the genome capacity of the AAV2 T=1 capsid.

## DISCUSSION

### Integration of molecular simulations, AI, and capsid theory as a framework for capsid design

Molecular simulations were a central component of our methodology, enabling assessment of the structural stability of capsid designs after mutations were introduced to resolve inter-subunit clashes. The field of molecular modeling of viral capsids has grown substantially since the first all-atom simulation of a capsid (Freddolino et al., 2006; Pérez-Segura et al., 2025; Perilla et al., 2016; Perilla & Schulten, 2017). When experimentally resolved structural templates are unavailable, capsid simulations often rely on idealized geometric or highly coarse-grained representations, such as patchy particles (Elrad & Hagan, 2008; Hagan & Chandler, 2006), trapezoidal particles (Rapaport, 2004, 2018), or simplified capsomer units. While these models are valuable for studying general principles of capsid assembly and mechanics, they sacrifice molecular-level detail and therefore cannot directly capture residue-specific interactions or interfacial rearrangements that are important for assessing capsid stability. This limitation motivates approaches that combine AI structure prediction, geometric capsid construction, and molecular dynamics to evaluate whether designed capsid architectures are physically stable at molecular resolution.

To contextualize the methodological leap of our approach, it is worth mentioning that molecular modeling and simulation techniques have long been used to investigate biomolecular dynamics (Schlick, 2010) when experimental methods are limited by resolution, cost, or accessibility. In virology, MD simulations have been widely applied to investigate the dynamics of viral structural components, including capsid and matrix proteins—for example, HIV-1 matrix–membrane interactions (Charlier et al., 2014; Monje-Galvan & Voth, 2020), structural flexibility of the hepatitis B virus capsid (Hadden et al., 2018), Ebola virus VP40 protein-membrane binding dynamics (Cioffi, Husby, et al., 2024; Cioffi, Sharma, et al., 2024; Pavadai et al., 2018), stability of CCMV capsid protein multimers (Szoverfi & Fejer, 2022), and calcium-dependent behavior of human metapneumovirus matrix protein (Leyrat et al., 2014)—often revealing mechanistic detail that is difficult or sometimes impossible to access experimentally. Additionally, simulations of entire viral capsids have been comparatively rare, largely due to the substantial computational expense and prior lack of suitable macromolecular modeling tools. Only a small handful of whole-capsid studies exist, typically based on experimentally determined structures from the Protein Data Bank (PDB) (Berman et al., 2000), including studies of HIV-1 capsid mechanics (Perilla & Schulten, 2017), HBV capsid function (Hadden & Perilla, 2018), and more recently ssDNA loaded AAV aggregation (Duran et al., 2026; Sharifi et al., 2025). Moreover, only a small number of full capsid structures are deposited in the PDB—representing a tiny fraction of the viruses known, and an even smaller fraction of the global virosphere, most of which remains uncultured. As a result, capsid-scale simulations remain far less accessible than simulations of smaller, resolved biomolecular systems.

Parallel advances in protein engineering have generated de novo icosahedral nanocompartments and protein shells (Hsia et al., 2016; Olshefsky et al., 2022), enabling applications in drug and gene delivery, vaccines, and immunotherapeutics (Bruun et al., 2018; Kanekiyo et al., 2013; Walls et al., 2020), enzyme encapsulation and nanoreactors (Azuma et al., 2018), structural scaffolds for materials science (Uchida et al., 2007), and orthogonal cellular compartmentalization/synthetic organelle engineering (Sigmund et al., 2018), among other applications (Hsia et al., 2016; Tetter et al., 2021). However, while these approaches demonstrate the power of de novo protein assembly, they do not directly address how to modify an existing viral capsid scaffold to form a larger, higher-order icosahedral architecture.

The use of AI-based folding methods, such as AlphaFold (Abramson et al., 2024; Jumper et al., 2021), and various other computational methods (Baek et al., 2021; Dauparas et al., 2022; R. Krishna et al., 2024; Lin et al., 2023; Watson et al., 2023) have made predicting, designing, and assembling oligomers from a given protein sequence an accessible task for researchers. Since the geometric theory of viral capsids identifies ideal tiles for building icosahedral capsids (Caspar & Klug, 1962; Twarock, 2004, 2005; Twarock & Luque, 2019), it provides a recipe for selecting optimal building blocks for capsid design directly from the protein sequence. In the case of AAV, the VP3 trimer was identified as the optimally folded oligomer, and its triangular shape, consistent with the dual hexagonal lattice, facilitated the initial construction of T=1 and T=3 capsids (Figure 2). However, the same block could be used to design T=4, T=7, or any larger icosahedral architecture. For other systems, the optimal oligomer compatible with the geometric theory of viral capsids may differ, but the same strategy used here can be applied to modify capsid architecture on other platforms.

### Uncertainties in the packaging capacity of the computationally designed T=3 capsids

The potential packaging capacity of the designed T=3 capsid was estimated from its internal volume. This is an upper-bound estimate of what the engineered shell could in principle enclose if the capsid remains stable under internal nucleic-acid load.

A further limitation is biological plausibility. AAV and related small icosahedral ssDNA viruses typically package genomes on the order of only a few kilobases, with AAV packaging approximately 4.7 kb (Hsi-Bell et al., 2025), and there is currently little evidence that conventional small icosahedral ssDNA virions package genomes far beyond this regime without major architectural changes (Mietzsch et al., 2025). The best-known larger ssDNA genome examples are generally associated with alternative virion organizations rather than a standard small icosahedral capsid, including the membrane-containing pseudo-icosahedral Finnlakevirus FLiP (∼9.2 kb) (Laanto et al., 2017), geminate particles in the family *Geminiviridae* (2.5–5.2 kb per genomic component) (Fiallo-Olivé et al., 2021), and the helical/coil-shaped archaeal virus ACV, which has an unusually large ∼24.9 kb ssDNA genome (Mochizuki et al., 2012; Prangishvili et al., 2020). For that reason, the upper end of the capacity estimated here is unlikely to be achieved by packaging a single canonical ssDNA molecule in a straightforward manner. It is more realistic to regard the highest values as theoretical volumetric maxima, although higher occupancies might become possible if multiple ssDNA molecules were packaged or if locally hybridized nucleic acid produced regions of dsDNA-like packing. Testing those possibilities would require explicit capsid–genome simulations and likely additional engineering of shell stabilization, which lies beyond the scope of the present study. Although full-capsid molecular simulations have advanced substantially, atomistic simulations of intact virions remain computationally expensive and are still limited to a relatively small number of systems, while genome-containing capsid simulations at near-complete scale have only recently become feasible in specialized studies (Coshic et al., 2024; Hadden & Perilla, 2018; Pérez-Segura et al., 2025). In that context, the coarse-grained approach used here is appropriate for evaluating capsid-scale structural integrity, global curvature, and long-timescale stability, even if it does not yet provide a direct test of the maximum realizable genome load. Thus, the packaging-capacity values reported here should be interpreted cautiously as structurally informed upper bounds rather than experimentally or computationally validated loading limits.

### Critical factors to investigate the assembly of AAV T=3 capsids experimentally

Ultimately, the most important next step is experimental validation. An important caveat to keep in mind is the fact that the AAV2 VP3Δ15 mutant containing deletions was able to form *in silico* a stable T=1 that may compete in the assembly process with T=3. Insertions could be strategically introduced to disfavor T=1 packing interactions—such as by modulating interface angles, local curvature constraints, or mechanical stress distributions that promote buckling or unfavorable internal pressure in the smaller architecture—while stabilizing the quasi-equivalent environments required for T=3 assembly. In this way, mutations would bias the energetic landscape toward the expanded capsid geometry rather than merely permitting both states. Experimental validation of these designs, including biochemical assembly assays and structural characterization, will be essential to determine whether selective T-number control can be achieved in practice and to assess the functional consequences of geometric expansion.

Regarding the possibility of repurposing existing AAV technologies to modify the tropism of T=3 capsids, it is worth noting that the surface properties, such as electrostatics and amino acid fluctuations (Figure S4), of the engineered T=3 are locally similar to those of T=1 (Figure S4). This is because T=3 contains the same local symmetrical protein regions as T=1. The only difference is in the additional quasi-equivalent positions that exist in the T=3 capsid, which generate a local 6-fold symmetry region that is not present in the T=1 capsid (Figure S4). This newly introduced region would be the primary area requiring careful assessment for altered interactions between a T=3 capsid and the immune system or host cell. At the same time, this local region could also be used as a protein binding target to adapt enzyme-linked immunosorbent assays (ELISA) to distinguish the presence of T=3 versus T=1 experimentally.

## CONCLUSION

In this work, we demonstrate that geometric principles of icosahedral symmetry can be leveraged to rationally expand the architecture of adeno-associated virus capsids beyond their native T=1 form. By integrating the geometric theory of capsids with AI-based structure prediction, procedural capsid construction, and coarse-grained molecular dynamics simulations, we generated a physically realistic T=3 AAV2 capsid model predicted to enclose an approximate fivefold increase in internal volume. Importantly, coarse-grained molecular dynamics simulations revealed that the engineered T=3 architecture remained structurally stable under physiological conditions, maintaining global integrity, curvature, and compactness throughout relaxation. This stability supports the geometric feasibility of T-number modulation and suggests that expanded AAV architectures can satisfy both symmetry constraints and mechanical requirements.

More broadly, this study establishes a generalizable computational framework for constructing and evaluating higher T-number capsids directly from sequence, including those lacking experimentally resolved structures. By coupling symmetry-guided design with physics-based simulation, our approach bridges schematic geometric models and dynamically relaxed capsid architectures, enabling quantitative assessment of stability, curvature compatibility, and capsid-scale mechanical behavior.

Although experimental validation is required to determine biological viability, these findings provide proof of principle that capsid size need not be treated as a fixed evolutionary constraint. Instead, T-number can be considered an engineerable parameter governed by geometric and energetic compatibility. Controlled manipulation of icosahedral architecture may therefore expand the functional landscape of viral vectors for gene therapy and broader biotechnological applications.

## METHODS

### Identifying optimal oligomeric tiles for icosahedral capsid design

The amino acid sequence of AAV2 VP3 was obtained from UniProt (P03135) (Bateman et al., 2025; Xie et al., 2002) and folded using AlphaFold3 (AF3) (Abramson et al., 2024). Oligomers ranging from one (n=1) to six (n=6) VP3 proteins were folded using multiple independent copies of the same sequence in AF3. Larger oligomers were beyond the permitted size in AF3 at the time of the study. The generated structures and associated pLDDT (predicted local distance difference test), pTM (predicted template modeling), and ipTM (interface predicted template modeling) scores were stored and compared. The optimal folded oligomer, VP3_n*_ (where n*=3), was identified as the one that yielded a high-quality score (>0.75). The symmetry and shape of the folded oligomers were assessed qualitatively. The qualitative polymeric shape of the optimal oligomer was compared to the shapes of the polyhedral tiles associated with the icosahedral lattices capable of forming homomeric capsids in the geometrical theory of icosahedral capsids: hexagonal (regular pentagons and hexagons), hexagonal-dual (regular triangles), trihexagonal-dual (rhombs), snub hexagonal-dual (florets), and rhombitrihexagonal-dual (kites) (Twarock & Luque, 2019).

### Building the VP3_3_ icosahedral capsids

The icosahedral lattice with a tile shape resembling that of the optimal oligomer was selected to design VP3_3_ icosahedral VP3_n_ capsids (Twarock & Luque, 2019). The folded structure of the optimal oligomer (triangular trimer, VP3_3_) was loaded into ChimeraX (Meng et al., 2023; Pettersen et al., 2021) and aligned to a single tile of a T(h=1,k=0) = 1 and a T(h=1,k=1) =3 icosahedral cage generated for the selected icosahedral lattice (hexagonal dual) using *hkcage* (Johnson & Speir, 1997). The cage radius was initially set to 130 Å to avoid clashing and get as close to the experimentally reconstructed AAV2 capsid (PDB ID: 1LP3) (Xie et al., 2002) as possible. The radius, R, for the T=3 cage was initially set according to the geometrical theory scaling, R(T) = R_0_·(T/T_0_)^1/2^, where T_0_ = 1 and R_0_=13 nm. The sphere factor was set to f = 0.9, sphere factors between 0.5 and 1.0 yielded similar results. The ChimeraX symmetry command, *sym* (Pettersen et al., 2021), was used to generate 19 and 59 additional copies of the triangular VP3 trimers (optimal oligomer), yielding the initial VP3_3_ T=1 and VP3_3_ T=3 capsid structures (see SI for *hkcage* and *sym* commands). Cage size was then refined based on visual inspection of trimer-trimer interface spacing after symmetrization.

### Coarse-graining protocol

The structural integrity and dynamic stability of T=1 and T=3 capsid architectures (AAV2 T=1, VP3_3_ T=1, VP3_3_ T=3, VP3_3_Δ15 T=3, VP3_3_Δ15 T=1) were evaluated using coarse-grained molecular dynamics (CGMD) simulations under physiological conditions. CGMD simulations are well suited for viral capsids because they reduce molecular complexity while retaining the dominant interactions needed to evaluate large-scale stability, curvature, elasticity, and inter-subunit organization. This is particularly useful for capsids, where key behaviors emerge from collective motions across large assemblies and timescales that are difficult to access with atomistic simulations. Coarse-grained models have therefore been widely used to study capsid assembly, mechanics, curvature, and stability across diverse viral systems (Grime et al., 2016; Hagan & Zandi, 2016; V. Krishna et al., 2010; Lynch et al., 2023). The capsid models were coarse-grained using the Martini 2.2P force field (Marrink et al., 2007; Monticelli et al., 2008). This force field incorporates an elastic network for proteins (ElNeDyn) (Periole et al., 2009), Martini 2.2 polar amino acids(de Jong et al., 2013), and polarizable water (Yesylevskyy et al., 2010). In this representation, amino-acids are coarse-grained to ∼5 Å resolution (Monticelli et al., 2008), with 3–5 heavy atoms mapped to each coarse-grained bead. This reduced resolution lowers computational cost while preserving key molecular interactions that contribute to capsid stability, including electrostatic and hydrophobic interactions. The elastic network was applied to each protein chain independently to maintain its folded three-dimensional structure. CHARMM-GUI (Brooks et al., 2009; Jo et al., 2008) webserver with the *Martini maker (Hsu et al., 2017; Qi et al., 2015)* plugin was used for generation of the AAV2 T=1, VP3_3_ T=1, VP3_3_Δ15 T=1. For the initial T=3 attempts and the VP3_3_Δ15 T=3 system, CHARMM-GUI (Brooks et al., 2009; Jo et al., 2008) with the *Martini maker (Hsu et al., 2017; Qi et al., 2015)* plugin was used for generating initial ‘include topology’ (.itp) files. The full VP3_3_Δ15 T=3 system, including the water box and salt, was created using the *insane* python script(Wassenaar et al., 2015). The capsids were solvated in polarizable Martini water and neutralized with 150 mM NaCl. The solvent box extended at least 12 Å from the capsid surface in all directions. Periodic boundary conditions were applied in all directions. The initial main system topology (.top) file was edited to reflect the actual system composition of protein chains, number of waters, and ions. The final sizes of each system are reported in the results section.

### Molecular dynamics simulations

Molecular dynamics simulations were performed to assess capsid stability and they were deployed with the usual three step protocol: *Energy Minimization*, *Equilibration*, and *Production*. All steps were completed using the Gromacs simulation software suite (Van Der Spoel et al., 2005).

#### Energy Minimization

The steepest descent algorithm slightly adjusted initial bead positions to remove minor unphysical overlaps, reduce inter-bead forces, and in turn lower the system’s potential energy. Energy minimization was carried out over a multi-step process to achieve a maximum force between any two CG beads of less than 500 kJ/mol/nm. This threshold follows common Gromacs and Martini practice, as Martini force fields employ relatively soft nonbonded potentials (σ ≈ 0.47 nm and є ≈ 2 − 6 kJ/mol) (Alessandri et al., 2019; Marrink et al., 2007) and therefore do not require all-atom-level minimization tolerances. A maximum force of 500-1000 kJ/mol/nm after energy minimization is widely used in Martini simulations because forces about ∼2000 kJ/mol/nm correspond to severe bead overlaps that produce accelerations large enough to destabilize the integrator at the 10-20 fs timesteps used for CG MD. In contrast, forces below ∼1000 kJ/mol/nm ensure that only mild steric compression remains, which is rapidly relaxed during the restrained equilibration phase. Larger thresholds (i.e., 10,000 kJ/mol/nm) indicate catastrophic geometric overlap and almost always lead to integrator instability and numerical blow-up. The process stopped when the maximum force on any bead was less than 500 kJ·mol^-1^·nm^-1^. For all five minimization steps, the initial step size was set to 0.005 nm. The output of each previous step was used as the input for the next. The tolerance for each respective step was 1000, 1000, 1000, 500, and 500 kJ·mol^-1^·nm^-1^. In the final minimization stage, the maximum force between any two beads was 377 kJ·mol^-1^·nm^-1^.

#### Equilibration

The system was then carefully equilibrated, also over a multi-step process, in an isothermal-isobaric (NPT) ensemble at 303.15 K and 1 bar. The isotropic pressure was maintained using a Berendsen barostat (Berendsen et al., 1984). The time constant for pressure coupling was set to 5.0 ps with a compressibility of 4.5·10^-5^ bar^-1^ in all directions, x, y, and z. The temperature was maintained using a velocity-rescaling thermostat (Bussi et al., 2007). The time coupling constant was 1.0 ps. Timesteps of 5 fs were used for all three equilibration steps. The total number of steps was 20,000 (0.1 ns) for both equilibration steps 1 and 2. The total number of steps was set to 160,000 (0.8 ns) for equilibration step 3. Total equilibration time was 1 ns. This amount of time was sufficient to achieve minimally fluctuating values around the target temperature and pressure.

#### Production

Production runs were performed in an NPT ensemble at the same temperature and pressure as equilibration using Gromacs (Van Der Spoel et al., 2005). The barostat was switched to the Parrinello-Rahman barostat (Parrinello & Rahman, 1981) to increase accuracy. The coupling constant was 12 ps. Long-range electrostatic interactions were approximated using reaction field electrostatics with an 11 Å cutoff applied to nonbonded interactions. The relative dielectric constant was set to 2.5; a standard value used in the Martini polarizable water model to ensure realistic dielectric behavior in hydrophobic regions while enabling water’s high dielectric constant (∼78.4) to emerge naturally from polarization effects (Yesylevskyy et al., 2010). Periodic boundary conditions were applied in x, y, and z directions. The leap-frog algorithm (Amini et al., 1987; Hockney et al., 1973) integrated Newton’s equations of motion with a timestep of 14 fs for a total of 2 µs (∼143,000,000 steps) for the WT T=1 and mutant T=3 systems. Timesteps were set to 20 fs for the T=1 PDB and T=1 mutant systems. Every 1 µs corresponds to roughly 3-8 µs of effective atomistic dynamics depending on the process and what motions dominate, since internal friction and solvent viscosity are reduced and the energy landscape is smoother (Marrink et al., 2004; Marrink & Tieleman, 2013; Monticelli et al., 2008). This timescale is sufficient to assess mechanical integrity since collective modes are sampled, like domain breathing, loop rearrangements, and local flexibility at interfaces. This timescale also exceeds those based on prior molecular dynamics studies of much smaller viral capsid systems (Freddolino et al., 2006; Hadden & Perilla, 2018) and is supported by relevant biophysical considerations, including the weak, collective nature of inter-subunit interactions and the timescales over which capsid assembly, rearrangement, and destabilization processes can emerge (Perkett & Hagan, 2014; Zlotnick, 2003). Frames were stored every 0.7 ns, yielding 2,000 frames for analysis. The frame rate and production time were set to minimize autocorrelations and enable an unbiased global stability analysis, based on prior studies (Arkhipov et al., 2006; Jana et al., 2023). The trajectories were visualized using Visual Molecular Dynamics (VMD)(Humphrey et al., 1996).

### Generating mutations compatible with the T=3 architecture

After capsid construction, multiple attempts to run MD failed during either the minimization or equilibration protocol due to atomic clashing. Some clashes were visually obvious; others were buried at the interface. Several rounds of minimization with multiple systems were performed with slight variations in the initial positions of the trimer subunits, which narrowed the problematic regions down to ∼15-20 amino acids, identified through careful visual screening. Hydrophobic residues in the interface were left unmodified to promote water expulsion. The resulting reduction in solvent-accessible hydrophobic surface area likely increases entropy through the release of ordered water molecules and may also enhance enthalpic stabilization through tighter van der Waals packing (Chandler, 2005; Dill, 1990; Lum et al., 1999; Pace et al., 1996; Tanford, 1978), collectively improving capsid stability. Hydrophobic contacts are collectively strong and resilient to changes in pH or ionic strength, providing more robust inter-subunit stabilization than geometry-dependent salt bridges or hydrogen bonds (Hendsch & Tidor, 1994; Horovitz et al., 1990; Jelesarov & Karshikoff, 2009; Serrano et al., 1990; Waldburger et al., 1995). Such conserved hydrophobic patches are a recurring feature of icosahedral capsid interfaces, often surrounded by polar or charged residues. Thus, the capsid was engineered such that the hydrophobic patches were conserved (and in some cases extended due to deletions) at the interface by removing patches of exposed residues that were both non-hydrophobic and non-faulty. Additionally, to avoid selecting potentially faulty mutants, published results on AAV2 capsids were reviewed to exclude VP3 mutations known to cause unstable capsids, inhibit capsid formation, reduce infectivity, or increase capsid disassembly, in which an exhaustive list is given by Wu et al. 2000 (P. Wu et al., 2000). After deletions were made, protein folding and T=3 capsid building process described for the WT VP3 protein in the initial section of the methods was repeated for the trimer containing the 15 deletions to assess if its multimeric structures were consistent with the geometrical theory of icosahedral capsids.

### Structural analysis and comparison of T=1 and T=3 capsids

The stability, packaging capacity, and molecular physicochemical properties of the engineered T=3 capsids were studied as described in the subsections below. Some of these properties were also analyzed for the VP3_3_ T=1 system as a control for VP3Δ15 T=3. Analyses of VP3Δ15 T=1 were carried to assess the impact of mutations on the canonical AAV2 capsid with respect to WT VP3 T=1. Additionally, because the Martini 2.2P coarse-grained model represents groups of atoms as beads with an effective length scale of approximately ∼0.50 nm (Monticelli et al., 2008), we included a conservative resolution-based uncertainty of ±0.25 nm for each averaged structural measurement displayed in the results. This term reflects the finite positional precision implied by the coarse-grained representation, rather than statistical sampling error alone.

#### Stability assessment

The global structural stability of a capsid was assessed by extracting the time evolution of the radius of gyration (*R*_g_) and radii components (R_avg_, R_min_, R_max_). All of the structural parameters were calculated with the Gromacs command line toolkit, using the compressed trajectory (.xtc), and the topology and coordinate files (.tpr and .gro), ChimeraX, or VMD. The radius of gyration (*R*_g_) reports on the global compactness of the capsid and was used to distinguish stable breathing-like fluctuations from large-scale expansion, collapse, or disassembly under the simulated conditions. To estimate stability time, each *R*_g_(*t*) trajectory was divided into sequential, non-overlapping 100 ns windows, allowing the local dynamical behavior of the trajectory to be evaluated over finite observation intervals (Cobo-López et al., 2023). Within each window, a local linear regression was fit to *R*_g_ as a function of time, where *m* is the fitted slope in units of nm/µs. The absolute slope, |m|, was used to quantify the magnitude of local structural drift in *R*_g_ within each 100 ns interval. A practical stability threshold of |m| < 0.25 nm/µs (half of model resolution) was applied; once a trajectory entered a window below this threshold and remained below it in all subsequent windows, the system was considered stable. The stability time was defined using the VP3_3_Δ15 T=3 trajectory because it was latest to enter a stable regime among the systems analyzed. At this time point, all other trajectories were also below the same threshold, providing a conservative common stability cutoff for subsequent analysis. The RMSD provides a complementary metric and was used to assess if the structure experienced any significant conformational changes, including collapse or spontaneous disassembly, with respect to its initial structural state (Figure S3). Together, these quantities provide a sufficient assessment of capsid stability under the simulated conditions.

#### Surface properties

The electrostatic surface calculations were analyzed to identify potential interaction hotspots on the capsid surface that may be critical for assembly, stability, or host interaction, and can help identify sites for charge-neutralization mutagenesis or antibody binding. The PDB structures were used to estimate the capsid and trimer Coulombic surface potential calculated chain-wise in ChimeraX (three monomers per trimer – averaged over all monomers gives us three sets of values). Coulombic surfaces can be seen in Figure S4. The root mean square fluctuations (RMSF) of backbone residues were calculated to provide residue-level insights into flexibility and dynamics, identifying regions that participated in inter-subunit contacts or dynamic interfaces versus those that remained structurally rigid—information that can guide mutagenesis experiments or epitope mapping. The RMSFs were calculated with the Gromacs command line toolkit and results are shown in Figure S4.

### Estimating capsid packaging capacity

The internal volume of the T=1 and T=3 capsids were calculated by using the Blob tool in ChimeraX(Meng et al., 2023), based on an average of five final frames of each simulation trajectory (1.90 µs, 1.92 µs, 1.94 µs, 1.96 µs, 1.98 µs). The resolution was set to 1.5 nm to adequately delineate the inner and outer capsid boundaries and density maps were generated. The inner capsid surface was selected with the Blob Picker tool which measured the enclosed volume. The total uncertainty in the average volume measurements was calculated by combining trajectory-sampling uncertainty and blob-resolution uncertainty in quadrature: 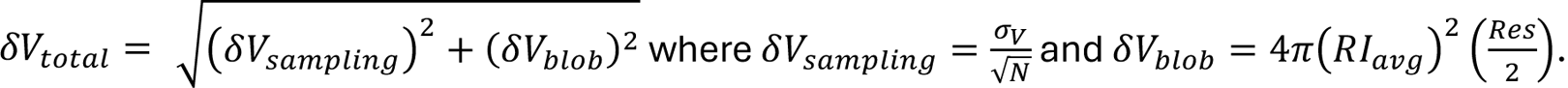 Trajectory sampling uncertainty was estimated as the standard error of the mean volume across sample frames. The blob-resolution uncertainty was estimated by multiplying the effective capsid surface area, approximated from the average inertial radius (RI_avg_) by half of the blob resolution (Res=1.5 nm). Inertial axes for the inner volume ellipsoid generated by the Blob tool were determined with the ChimeraX command *measure inertia*. The results for interior volume were then displayed as: V_int_ (±δV). The error in volume mostly comes from the resolution of the Blob probe, which needed to be large enough to sufficiently close all “holes” in the capsid surface. Using these volume measurements, we were then able to compute an estimated genome packaging capacity for the simulated VP3Δ15_3_ T=3 capsid.

Wild-type AAV2 was treated as having a single-stranded DNA genome size (G±δG) of approximately 4.8 ± 0.2 kb, consistent with the 4,679-bp AAV2 reference genome (NCBI RefSeq NC_001401.2) and the commonly reported ∼4.7–5.0 kb AAV genome/package-size range, depending on the specific reference sequence, whether the value is reported as the exact WT genome or rounded packaging capacity, and whether recombinant vector elements are included (Adachi et al., 2015; Dong et al., 1996; Kosaka et al., 2025; Z. Wu et al., 2010). Using this genome size (G±δG) and the measured V (±δV) of the simulated AAV2 T=1 capsid, the estimated packaging density (ρ±δρ) of the genome was calculated with ρ±δρ=(G±δG) / (V±δV). Then, from ρ±δρ and the measured V±δV of the simulated VP3Δ15_3_ T=3 capsid, we estimated the total genome capacity (G±δG) using (ρ±δρ)* (V±δV)=G±δG. The value for G_max_ was then taken as the upper bound of the uncertainty in G±δG value for AAV2 T=1 and VP3Δ15_3_ T=3.

Uncertainties for all calculated quantities involving multiplication or division were propagated using standard first-order error propagation. For products and ratios, the propagated relative uncertainty was calculated as the quadrature sum of the relative uncertainties of the input quantities:

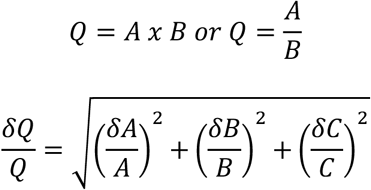

The absolute uncertainty was then obtained by multiplying the calculated central value by the propagated relative uncertainty:

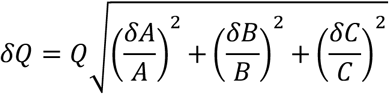

Final values were reported as symmetric uncertainties in the form Q±δQ, where Q was calculated from the nominal central values.

### Hardware

Minimization, equilibration, production, and analysis with the Gromacs package were carried out on a Dell Precision 7960 Tower workstation containing a 56-core, 112-thread Intel Xeon w9-3495X processor, dual Nvidia RTX 5000 Ada Generation GPUs, and 256 GB of DDR5 RDIMM ECC memory. Visualizations of capsids and trajectories were obtained on a similar workstation with four times more memory, 1 TB of DDR5 RDIMM ECC memory, and dual Nvidia RTX 5000 Ada Generation GPUs.

## ACKNOWLEDGEMENTS

AL thanks Matt Shtrahman (University of California, San Diego) for an inspiring discussion about AAV at a San Diego Microbiology Group (SDMG) meeting in 2019. AL also thanks Pantelis Tsoulfas, Ronald Desrosiers, Cynthia Silveira, Imran Noor, and Ian Fitzmaurice at the University of Miami for their insights. AL acknowledges support from the National Science Foundation (Award #2424579) and internal support from the University of Miami through the 2025 Provost Award and the Sylvester - College of Arts and Sciences - Catalysts for Cure Program. AL and MC acknowledge support from the Gordon Betty Moore Foundation (Award GBMF9871, https://doi.org/10.37807/GBMF9871).

