## Supplementary Information for "Computational Rational Design of Larger AAV Icosahedral Capsids"

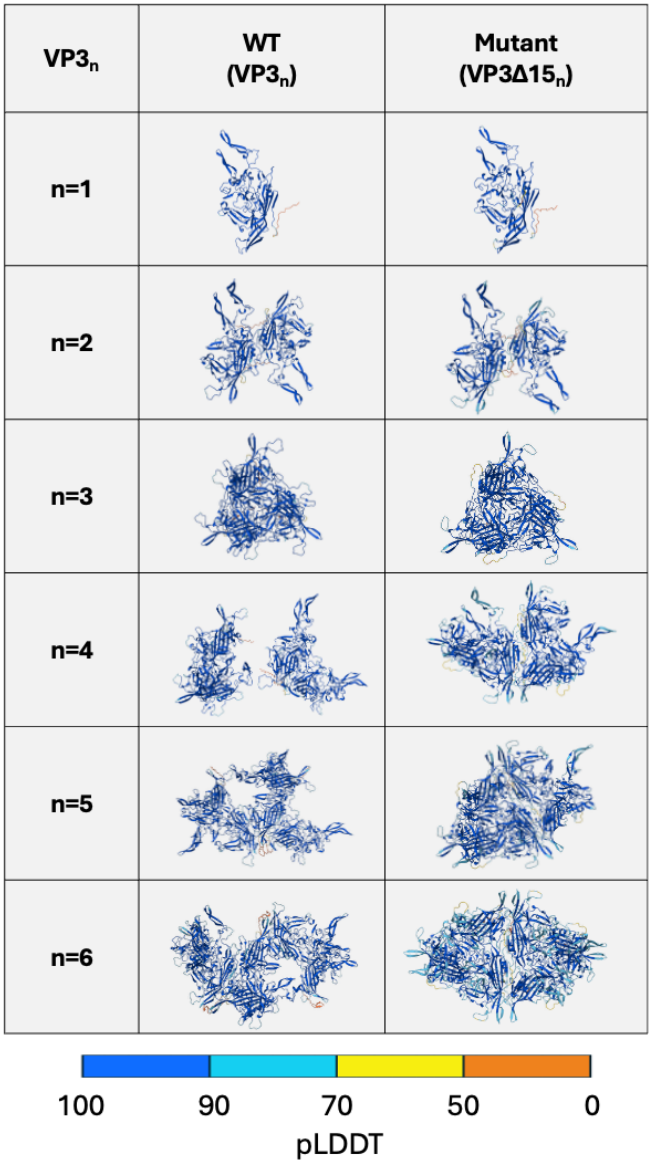


**Figure S1. Multimer analysis for VP3$_n$ and VP3$_n$Δ15.** AlphaFold 3 results for the folded AAV2 wildtype and mutant VP3 monomer and oligomers built by up to six subunits (n). A predicted local distance difference test (pLDDT) scale bar is shown below the folded oligomers.

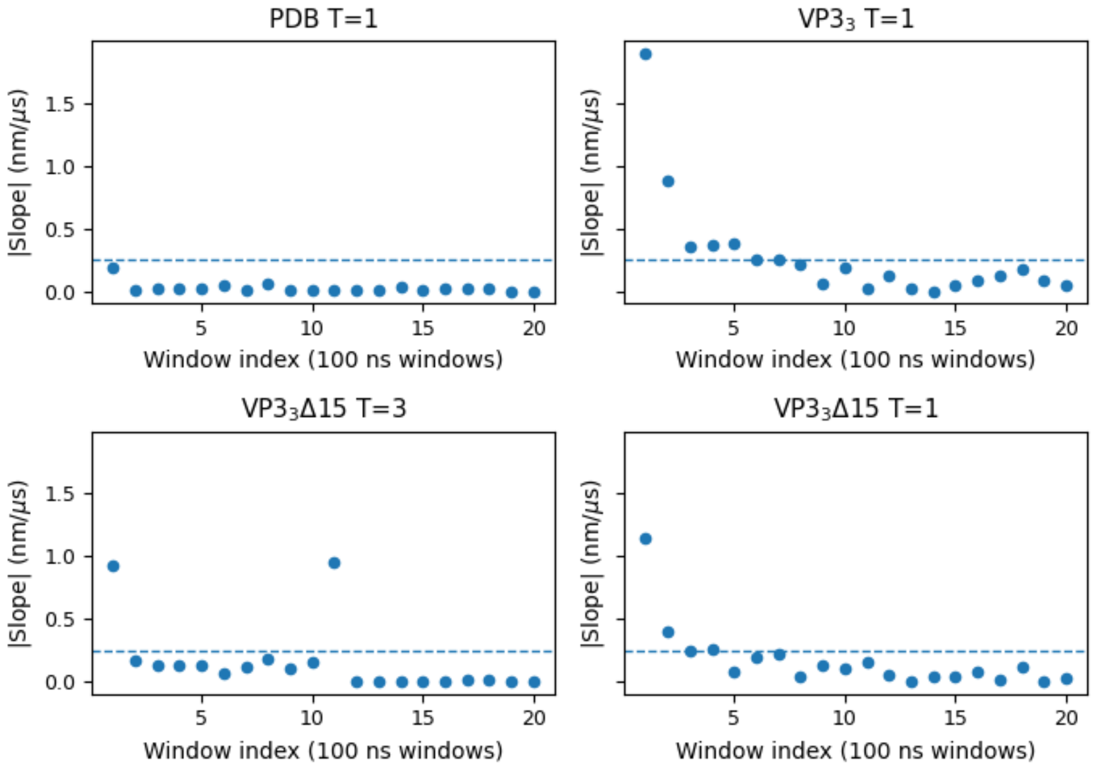


**Figure S2. Block-wise assessment of structural stability over 100 ns intervals for all systems.** Slope values are plotted for each 100 ns block for each system, where each data point is located at the end of each block (e.g., the slope value of 0-0.1 μs is located at window index 1). The coarse-grained resolution was used as a slope threshold to distinguish between change and no change where it is displayed by a dashed line at 0.25 nm (model resolution).

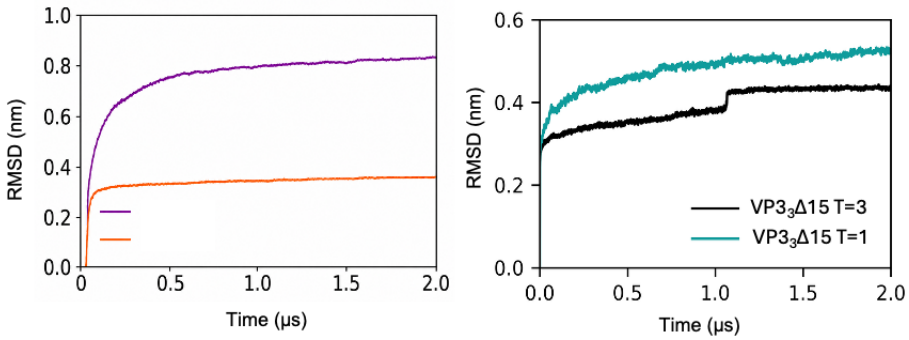


**Figure S3. Time evolution of the root mean square deviations.** The root mean square deviation (RMSD) was calculated per frame for each system (AAV2 PDB T=1, AAV2 VP33 T=1, AAV2 VP3Δ15$_3$ T=1, AAV2 VP3Δ15$_3$ T=3). The system coordinates obtained post-equilibration (pre-production) were chosen as the refence structure.

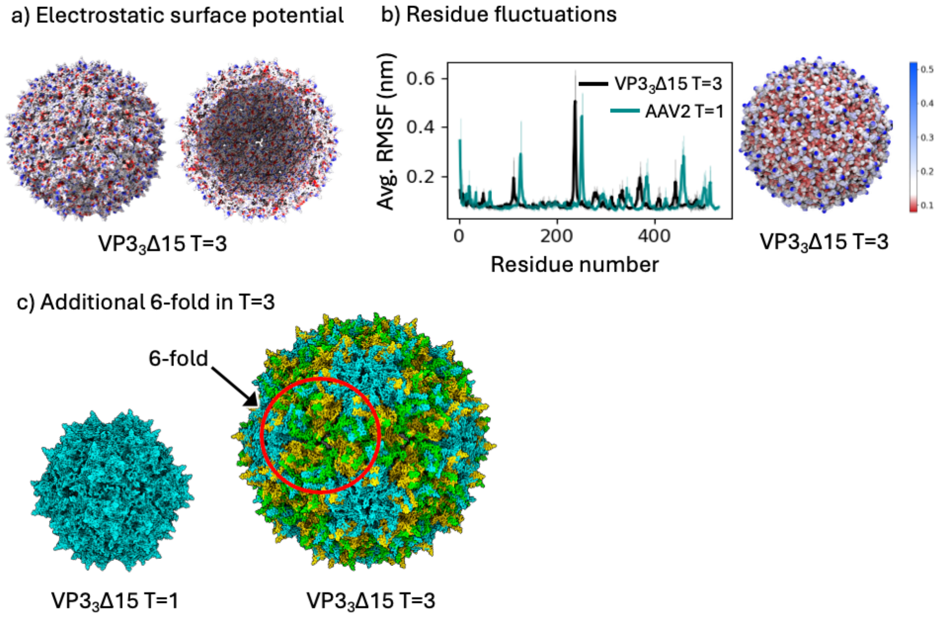


**Figure S4. Surface properties of VP3$_3$Δ15 T=3, AAV2 T=1, and VP3$_3$Δ15 T=1.** (a) Electrostatic surface map of VP3$_3$Δ15 T=3. The Coulombic surface coloring feature was applied to highlight regions of positive and negative charge, providing insight into the electrostatic landscape of the capsid. (b) The root mean square fluctuations (RMSF) of the backbone (BB) beads of the capsid averaged over all monomers for $VP3_3\Delta15$ T=3 and AAV2 T=1. (c) The additional quasi-equivalent position that exists in the T=3 capsid, which generates a local 6-fold symmetry region that is not present in the T=1 capsid.

**Table S1. Residues avoided when selecting deletions.** Amino acid residues were avoided if they were hydrophobic and/or if they appeared in Wu et al. 2000 as faulty (unstable, disassembly, or no capsid formation).

| Target region | Avoided (hydrophobicity) | Avoided (Wu et al. 2000) |
|---|---|---|
| N656-A667 | P657, T659, F661, A663-A667 | A664, K665 |
| N695-G718 | N695-S702, V708, V710, F712, V714, G718 | X |
